# A murine model of the human CREBRF^R457Q^ obesity-risk variant does not influence energy or glucose homeostasis in response to nutritional stress

**DOI:** 10.1101/2021.05.06.442909

**Authors:** Jitendra S. Kanshana, Polly E. Mattila, Michael C. Ewing, Ashlee N. Wood, Gabriele Schoiswohl, Anna C. Meyer, Aneta Kowalski, Samantha L. Rosenthal, Sebastien Gingras, Brett A. Kaufman, Ray Lu, Daniel E. Weeks, Stephen T. McGarvey, Ryan L. Minster, Nicola L. Hawley, Erin E. Kershaw

## Abstract

Obesity and diabetes have strong heritable components, yet the genetic contributions to these diseases remain largely unexplained. In humans, a missense variant in Creb3 regulatory factor (CREBRF) [rs373863828 (p.Arg457Gln); CREBRF^R457Q^] is strongly associated with increased odds of obesity but decreased odds of diabetes. Although virtually nothing is known about CREBRF’s mechanism of action, emerging evidence implicates it in the adaptive transcriptional response to nutritional stress downstream of TORC1. The objectives of this study were to generate a murine model with knockin of the orthologous variant in mice (CREBRF^R458Q^) and to test the hypothesis that this CREBRF variant promotes obesity and protects against diabetes by regulating energy and glucose homeostasis downstream of TORC1. To test this hypothesis, we performed extensive phenotypic analysis of CREBRF^R458Q^ knockin mice at baseline and in response to acute (fasting/refeeding), chronic (low- and high-fat diet feeding), and extreme (prolonged fasting) nutritional stress as well as with pharmacological TORC1 inhibition. The results demonstrate that the murine CREBRF^R458Q^ model of the human CREBRF^R457Q^ variant does not influence energy/glucose homeostasis in response to these interventions. Alternative preclinical models and/or studies in humans will be required to decipher the mechanisms linking this variant to human health and disease.

## Introduction

Obesity is a global public health threat that is associated with additional metabolic abnormalities (i.e., insulin resistance, dyslipidemia) that increase the risk of common diseases such as diabetes and cardiovascular disease. Although obesity clearly has a complex and multifactorial etiology, it has a strong heritable component (>80% in some studies). Over the past several decades, large genome-wide association studies (GWAS) have sought to identify the genetic contributions to this heritability. These studies have been instrumental in linking numerous genes/loci to anthropometric and/or metabolic traits, including body mass index (BMI)/obesity (1, 2), regional adiposity/insulin biology (2, 3), and glycemia/diabetes (4). These studies have revealed important insights into the underlying physiology/pathophysiology of these traits such as the critical role of central neuroendocrine regulation of energy homeostasis in obesity (1, 2), adipose tissue biology in insulin resistance (2, 3), and beta cell biology in diabetes (4). Despite these insights, the majority of heritability of these metabolic traits remains largely unexplained. For example, genes/loci identified by GWAS account for less than 3% of the variance in BMI (1). Furthermore, these GWAS are historically enriched in populations with European ancestry and have not adequately assessed genetic contributions from underrepresented groups (5), many of which are disproportionately affected by obesity, diabetes, and cardiometabolic diseases. Identifying novel genes/pathways underlying this “missing heritability” across diverse populations could improve prevention and/or treatment of obesity and its complications.

To bridge this knowledge gap, our research group has established an ongoing collaboration with the Samoan Ministry of Health to understand and address cardiometabolic disease risk in Samoans, a geographically isolated population with a particularly high prevalence of obesity. This collaborative effort resulted in the identification of a missense variant in Creb3 regulatory factor (CREBRF) [rs373863828 (p.Arg457Gln); CREBRF^R457Q^] by GWAS in Samoans that was strongly associated with BMI (6). This obesity-risk variant (odds ratio for obesity of 1.305 per copy) is unique in that, even though it has a BMI-increasing effect size greater than any other known common obesity-risk variant (1.36–1.45 kg/m^2^ CREBRF^R457Q^ versus only 0.37 kg/m^2^ for the FTO locus), it is paradoxically associated with relative protection from diabetes (odds ratio for diabetes of 0.586 after controlling for BMI) and other metabolic complications of obesity (7). This variant is also positively related to linear height in humans (8-10). Although this variant is extremely rare in populations most commonly studied in GWAS (i.e., Europeans), it is very common in Samoans (∼46% carry at least one copy of the risk allele). The relationship between CREBRF^R457Q^ and obesity/diabetes risk has now been replicated by numerous other groups in Oceanic populations (9, 11-15). However, despite its large effect size and high prevalence in this population, the mechanisms by which the CREBRF^R457Q^ variant contributes to obesity and obesity-associated traits remain unknown.

CREBRF (also known as Luman/CREB3 Recruitment Factor, LRF) was initially identified in 2008 as a novel protein that binds to CREB3 and inhibits CREB3-mediated activation of the unfolded protein response (UPR) in mice (16). Global targeted deletion of CREBRF in mice results in lower total body mass (17), suggesting that the human CREBRF^R457Q^ variant may be a gain-of-function variant with respect to body mass / energy homeostasis. This hypothesis is further supported by our data demonstrating that ectopic overexpression of the human CREBRF^R457Q^ variant in a murine 3T3-L1 adipocyte model promotes adipogenesis, decreases mitochondrial respiration, and increases fat storage (6). While these data implicate adipocytes in the pathophysiology of the CREBRF^R457Q^ variant, the relative contributions of other tissues and cell types where CREBRF is also expressed remain unknown (6). CREBRF is highly regulated by nutritional status and TORC1 action (increased by fasting/starvation and TORC1 inhibition; decreased by refeeding and TORC1 activation) (6, 18). Overexpression of CREBRF in 3T3-L1 adipocytes protects against starvation (6), whereas deletion of CREBRF in Drosophila S6 cells, larvae, and adults increases susceptibility to starvation (18). These data suggest that CREBRF plays a critical role in the cellular and physiological response to nutritional stress. In addition, mechanistic studies in Drosophila demonstrate that CREBRF mediates the majority of the downstream effects of TORC1 inhibition by rapamycin (18). Although similar findings have yet to be confirmed in mammals, these data nonetheless implicate CREBRF in one of the most well-known energy sensing pathways in biology.

Despite the potential importance of these findings, virtually nothing is known about the function or physiological relevance of CREBRF or the CREBRF^R457Q^ variant. Our overall goal is to understand how the CREBRF^R457Q^ variant influences energy and metabolic homeostasis. Based on the above data, we hypothesized that the CREBRF^R457Q^ variant promotes obesity and protects against diabetes by regulating gene pathways involved in energy and glucose homeostasis downstream of TORC1. To test this hypothesis, we generated an animal model of the human CREBRF^R457Q^ variant by using CRISPR-Cas9 gene-editing technology to knockin the orthologous mutation in mice (CREBRF^R458Q^). We then phenotyped these mice at baseline and in response to acute (fasting/refeeding), chronic (low- and high-fat diet feeding), and extreme (prolonged fasting) nutritional stress as well as with pharmacological TORC1 inhibition.

## Materials and Methods

### Generation and validation of *Crebrf* knockin (*Crebrf*KI) mice

CREBRF^R458Q^ knockin (*Crebrf*KI) mice were generated using CRISPR-Cas9 gene-editing technology in a congenic C57BL/6J mouse background strain with the assistance of the University of Pittsburgh Transgenic and Gene Targeting (TGT) Core (19-21). Briefly, a double-strand DNA break was introduced in proximity of the murine R458 codon by targeting the Cas9 nuclease with a small guide RNA (sgRNA) (murine *Crebrf* target sequence: 5’-CTGGTACATATTACTTGGCA-3’, chr17:26,757,961-26,757,980 (mm10)). The desired genomic DNA substitutions were introduced by co-injection of a PAGE-purified single stranded oligodeoxynucleotide (ssODN) ultramer to serve as template for homology-directed repair (HDR). Two substitutions (lower cap bold in the sequence below; see also **Fig 1A**) were introduced: i) a silent substitution (TTC to TcC) to introduce a BamH1 site (underlined) approximately 20 bp away from the R458 codon, to serve as a proxy for the mutation for genotyping purposes; and ii) a coding substitution (CGA to CaA) to introduce the desired R458Q knockin variant. A PCR-generated sgRNA template was used for sgRNA synthesis using the MEGAshortscript T7 Kit (ThermoFisher Scientific), and Cas9 mRNA was produced using a linearized plasmid as template for the in vitro transcription mMESSAGE mMACHINE T7 Ultra Kit (ThermoFisher Scientific). Both sgRNA and Cas9 mRNA were purified using the MEGAclear Kit (ThermoFisher Scientific). C57BL/6J pronuclear-stage zygotes, obtained by natural mating of superovulated females, were microinjected with sgRNA (10 ng/μL, each) and Cas9 mRNA (20 ng/μL), and the ssODN (1µM). Injected zygotes were cultured overnight and 2-cell embryos were transferred to pseudopregnant CD1 mice to obtain potential founder mice.

**Fig 1.**
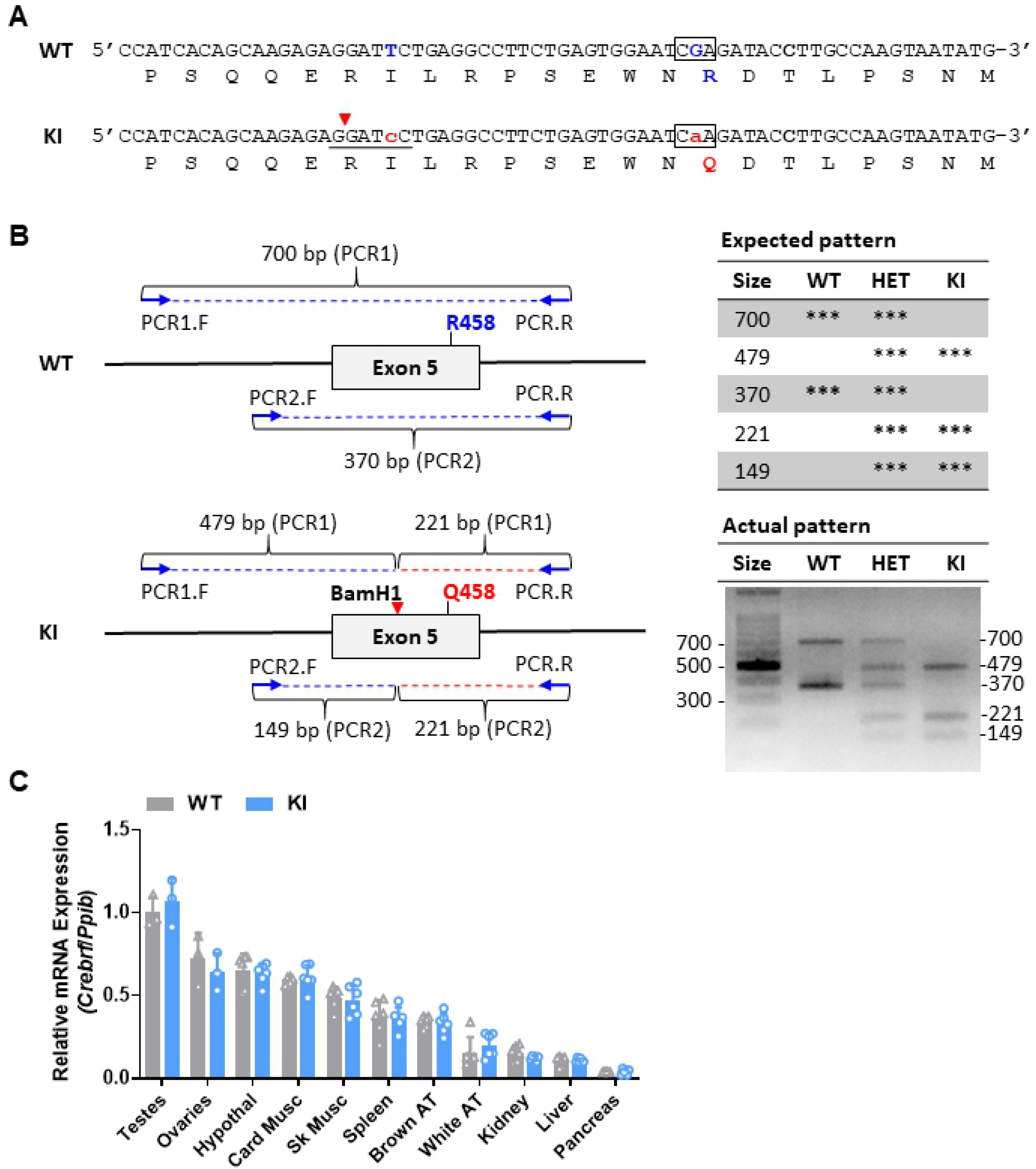
Generation and validation of *Crebrf* knockin (*Crebrf*KI) mice. **(A)** The CREBRF^R458Q^ knockin allele was generate using CRISPR-Cas9 gene-editing technology. Genomic (top) and protein (bottom) sequence of the wild-type (WT) and gene-edited knockin (KI) alleles are shown. To generate the KI allele, two substitutions were introduced: i) a silent substitution (TTC to TcC) to introduce a BamH1 site (underlined with restriction cut site shown with red arrowhead) about 20 bp away from R458 codon, to serve as a proxy for the mutation for genotyping purpose; and ii) a coding substitution (CGA to CaA) to introduce the desired R458Q knockin variant (surrounded by the black box); **(B)** The WT and KI alleles were confirmed by direct sequencing and restriction length polymorphism (RFLP) analysis. The schematic for the two-step nested PCR RFLP assay is shown on the left, and the resulting representative gels, confirming homozygous WT, heterozygous (Het), and homozygous KI mice, are shown on the right. **(C)** *Crebrf* mRNA expression by qPCR in multiple tissues of WT and KI mice (M and F, 6-week-old, ad lib fed LFD, n=6/group except testes and ovaries n=3/group). Data are murine *Crebrf* relative to reference gene *Ppib* and expressed relative to WT testes. Testes, ovaries, hypothalamus (hypothal), cardiac muscle (Card Musc), gastrocnemius-plantaris-soleus skeletal muscle (Sk Musc), spleen, brown adipose tissue (brown AT), perigonadal white adipose tissue (white AT), liver, and pancreas show no genotype effects on Crebrf mRNA expression. Unpaired two-tailed t-tests were used to compare gene expression (dependent variable) between genotype (fixed factor). Data are mean (SD). p<0.05 was considered significant.

The presence or absence of the desired knockin allele was confirmed in several ways. First, purified tail DNA from potential founders were screened using a two-step nested PCR-RFLP assay. The first amplification (with genomic DNA as template) used forward primer 5’ TTTAATGCCTGGCACCATTT and reverse primer 5’ GAACGAGGCAGAGGATTCAA to generate a 700 bp product (PCR product 1, PCR1). The second amplification (with PCR product 1 as template) used forward primer 5’ TGACAATTGTGGGACCATGT and reverse primer 5’ GAACGAGGCAGAGGATTCAA to generate a 370 bp product (PCR product 2, PCR2). Both steps used PCR conditions of 95°C x 30s / 55°C x 30s / 72°C x 30s for 35 cycles. The second PCR product was then digested with BamH1 and resolved on agarose, resulting in 370 bp (plus 700 bp) bands for the wild-type (WT) allele and a 221 bp and 149 bp (plus 479 bp) bands for the knockin (KI) allele (**Fig 1B**). Second, for founders positive for the KI allele by the PCR-RFLP assay, the 700 bp PCR products were subcloned into the pCR™4-TOPO^®^ TA vector and transfected into *Escherichia coli* DH5α cells (TOPO 4 Kit, ThermoFisher Scientific) followed by plasmid preparation and direct Sanger sequencing. In this way, a founder carrying the desired KI allele was confirmed and then mated to WT C57BL/6J mice to expand the colony. Subsequently, mice were genotyped using our previously published locked nucleic acid assays (21). All oligonucleotide sequences used in this study are listed in **S1 Table**.

### Animals

Mice were housed under standard conditions (25°C, 14:10 h light:dark cycle) with *ad libitum* access to a control low-fat diet (LFD: Research Diets D12450Hi; 10/70/20 kcal% fat/carbohydrate/protein; 3.82 kcal/g) or high-fat diet (HFD: Research Diets D12451i; 45/35/20 kcal% fat/carbohydrate/protein; 4.73 kcal/g) from weaning until sacrifice, except as otherwise noted (i.e., during fasting). The LFD was specifically formulated as the control for the HFD. For the longitudinal comprehensive phenotyping experiment, male and female WT and KI mice were fed LFD and HFD until 22 weeks of age and then sacrificed in fasted (16h) or refed (24h of fasting followed by 16h of refeeding) states. For the prolonged fasting experiment, a separate cohort of female WT and KI mice were ad lib fed HFD as above until 36 weeks of age. After acclimation to single housing in metabolic cages (Sable Promethion system), they were then fasted until they lost 25% of their initial body weight and then refed until they re-established their initial body weight. For in vivo mTORC1 inhibition experiments, a separate cohort of male WT and KI mice were ad lib fed LFD or HFD as above, but were treated with either vehicle (Veh: sterile 10% PEG 400/8% ethanol plus an equal volume of 10% Tween 80) or rapamycin (Rapa: rapamycin 4 mg/kg body weight dissolved in the above vehicle; rapamycin was obtained LC Laboratories) via an intraperitoneal (ip) injection every other day for three weeks prior to sacrifice in the ad lib fed state at 22 weeks of age based on a previously reported protocol (22). Animal experiments were approved by the University of Pittsburgh IACUC (Protocol #20107971) and conducted in conformity with PHS Policy for Care and Use of Laboratory Animals (PHS Assurance #D16-00118).

### Metabolic and biochemical analyses

Body composition, energy expenditure, and metabolic parameters were performed as described (23-25). Body composition was determined by echo magnetic resonance imaging (Echo Medical Systems). Energy intake and expenditure, energy substrate utilization, and activity were determined by indirect calorimetry (Promethion Metabolic Phenotyping Cages from Sable Systems International). Plasma glucose was measured using a One-Touch FastTake glucometer (Lifescan). For glucose tolerance tests (GTTs), mice were injected ip with 1.5 g/kg glucose following a 6-h fast. Other serum parameters were determined using the following kits: insulin (Ultra Sensitive Mouse Insulin ELISA kit; Crystal Chem), triacylglycerol or TAG (Pointe Scientific Triglycerides Liquid Reagents; Fisher Scientific); non-esterified fatty acids or NEFAs (HR series NEFA-HR[2] Reagents; Wako Diagnostics), leptin (mouse leptin ELISA, Crystal Chem); and adiponectin (mouse adiponectin ELISA, Crystal Chem). Liver TAGs (Infinity Triglycerides Liquid Stable Reagent; Thermo Scientific) were determined following tissue lipid extraction in chloroform:methanol (2:1), followed by acidification with H_2_SO_4_ and phase separation by centrifugation, and reconstitution in 60 μl of tert-butanol plus 40 μl of a 2:1 mixture of Triton X-114 (23).

### Gene expression analysis

RNA was isolated using RNeasy Lipid Tissue Mini Kit with on-column DNase treatment (Qiagen) and then reverse transcribed into cDNA using qScript Supermix (Quanta Biosciences). Relative gene expression was then determined by quantitative PCR (qPCR; ABI QuantStudio3 System) using the standard curve method after amplification of cDNA with gene-specific primer-probe sets using PerfeCTa Fastmix II Fastmix (Quantabio). Taqman Gene Expression Assays were Mm01299053_m1 for *Crebrf* and Mm02342430_g1 for reference gene *Ppia* (ThermoFisher). qPCR was conducted in accordance with MIQE guidelines (26).

### Protein analysis by immunoblotting

All antibodies used in this study are listed in **S2 Table**. Immunoblotting was performed as described (27). Briefly, tissues were homogenized in ice-cold RIPA lysis buffer (250mM Tris-HCl, pH 7.4, 750 mM NaCl, 5% Triton X-100, 2.5% sodium deoxycholate, 0.5% sodium dodecyl sulphate (SDS), 100 mM NaF, 2 mM Na_3_VO_4_, 1 mM phenylmethylsulfonyl (PMSF) and 1% cocktail protein protease and phosphatase inhibitors (Sigma), pH 8.0). Lysates were centrifuged at 14,000x *g* for 30 min at 4 °C, and supernatants were stored at -80 °C until analysis. Lysate protein was resolved by SDS-PAGE (Novex NuPAGE 4-12% Bis-Tris gels, Invitrogen) and transferred to polyvinylidene difluoride (PVDF) membranes (Millipore sigma). Phosphorylated and total proteins were identified by immunoblotting using the following primary antibodies: Phospho-S6 Ribosomal Protein (Ser240/244) Antibody #2215 and α-Tubulin (DM1A) Mouse mAb #3873 (Cell Signaling Technologies) at 1:1000 dilution. Immunoblots were evaluated using IRDye® 680RD donkey anti-mouse IgG and 800CW donkey anti-rabbit IgG (LI-COR) at 1:15,000 and scanned on an Odyssey Clx Imaging System (LI-COR). Fluorescent signals were quantified using Image Studio Software (LI-COR).

### Statistical analysis

Results are expressed as mean (SD). Comparisons were made using general linear models followed by determination of simple effects for pair-wise comparisons if relevant (SPSS Software v27). For repeated measurements, comparisons were made by generalized linear model with time as a within subject variable. For indirect calorimetry data, comparisons were made using generalized estimated equations with time as a within subject variable. Dependent variables and fixed effects for each analysis are indicated in the text and figure legends. Males and females were analyzed separately. For all analyses, p values of <0.05 were considered statistically significant.

### Data and resources availability

1) Data sharing. Data supporting the findings, methods, and conclusions in this manuscript are available within the manuscript in the supplemental online material, or from the corresponding author. 2) Resource sharing. Information related to antibodies, primers, and key reagents are summarized in supplemental online material. Unique resources used in this manuscript are available from corresponding author. The *Crebrf*KI mouse model or its derivatives (embryos, sperm) will be available for distribution from the corresponding author or deposited with a commercial vendor.

## Results

### Generation and validation of *Crebrf* knockin (*Crebrf*KI) mice

Our previous study identified a nonsynonymous variant in CREBRF (rs373863828) to be strongly associated with obesity and diabetes risk in humans (6). The variant is a guanine (G) to adenine (A) substitution that changes the amino acid at position 457 from an arginine [Arg, R] to a glutamine (Gln, Q), designated CREBRF^R457Q^ in humans (**S1A-B Fig**). CREBRF DNA and protein sequences are highly conserved in vertebrates. Comparison of human to mouse protein sequences revealed that the homologous amino acid in mice was at position 458 (**S1B Fig**). To generate *Crebrf*KI mice, CRISPR-Cas9 gene-editing technology was used to replace the guanine (G) to adenine (A) in genomic DNA of congenic C57BL/6J mice (comparable to the rs373863828[A] minor allele in humans), to generate the homologous mutant protein, designated CREBRF^R458Q^ in mice. The annotated murine wild-type (WT) and genome-edited knockin (KI) alleles (**Fig 1A**) were confirmed by direct sequencing and restriction fragment length polymorphisms (RFLP) (**Fig 1B**). Resulting mice homozygous for the KI allele are subsequently designated *Crebrf*KI or KI mice. Gene expression analysis confirmed *Crebrf* mRNA expression across multiple tissues in WT and KI mice, and this expression did not differ between genotypes (**Fig 1C**). Endogenous CREBRF protein expression is known to be low and transiently expressed at baseline (16). Consistently, CREBRF protein was below the limit of detection in peripheral tissues of both WT and KI mice. Experimental WT and KI mice, generated from heterozygous x heterozygous crosses, were viable and fertile with the expected Mendelian ratios of genotype and sex in resulting pups.

### The CREBRF^R458Q^ variant does not affect body weight, body composition, or linear growth at baseline (low-fat diet) or in response to chronic nutritional overload (high-fat diet) in mice

The human CREBRF^R457Q^ variant is associated with greater body mass (6, 11-13) and linear height (8-10) in humans. To determine the effect of the murine CREBRF^R458Q^ variant on these parameters, we evaluated energy homeostasis, body composition, and linear growth in male and female WT and KI mice under control nutritional conditions (low-fat diet, LFD; males only) and in response to chronic nutritional overload (high-fat diet, HFD; males and females) from weaning until sacrifice at 22 weeks of age. No differences were identified between genotypes for longitudinal total body mass (**Fig 2A**), fat mass (**Fig 2B**), or lean mass (**Fig 2C**) in male (**Fig 2, blue, top row**) or female (**Fig 2, red, bottom row**) mice on either diet. In addition, at the time of sacrifice, no differences were identified between genotypes for two different measures of linear growth: tibia length and nose-rump length (**Fig 2D**) or for individual tissue weights (**S2A-B Fig**). Notably, as positive controls, the expected effects of diet on these parameters were present, i.e., higher total body mass and fat mass with HFD-feeding (**Fig 2A-B
top row**; **S2A Fig**). Thus, the murine CREBRF^R458Q^ variant does not affect body weight, body composition, or linear growth at baseline or in response to chronic nutritional overload in mice.

**Fig 2.**
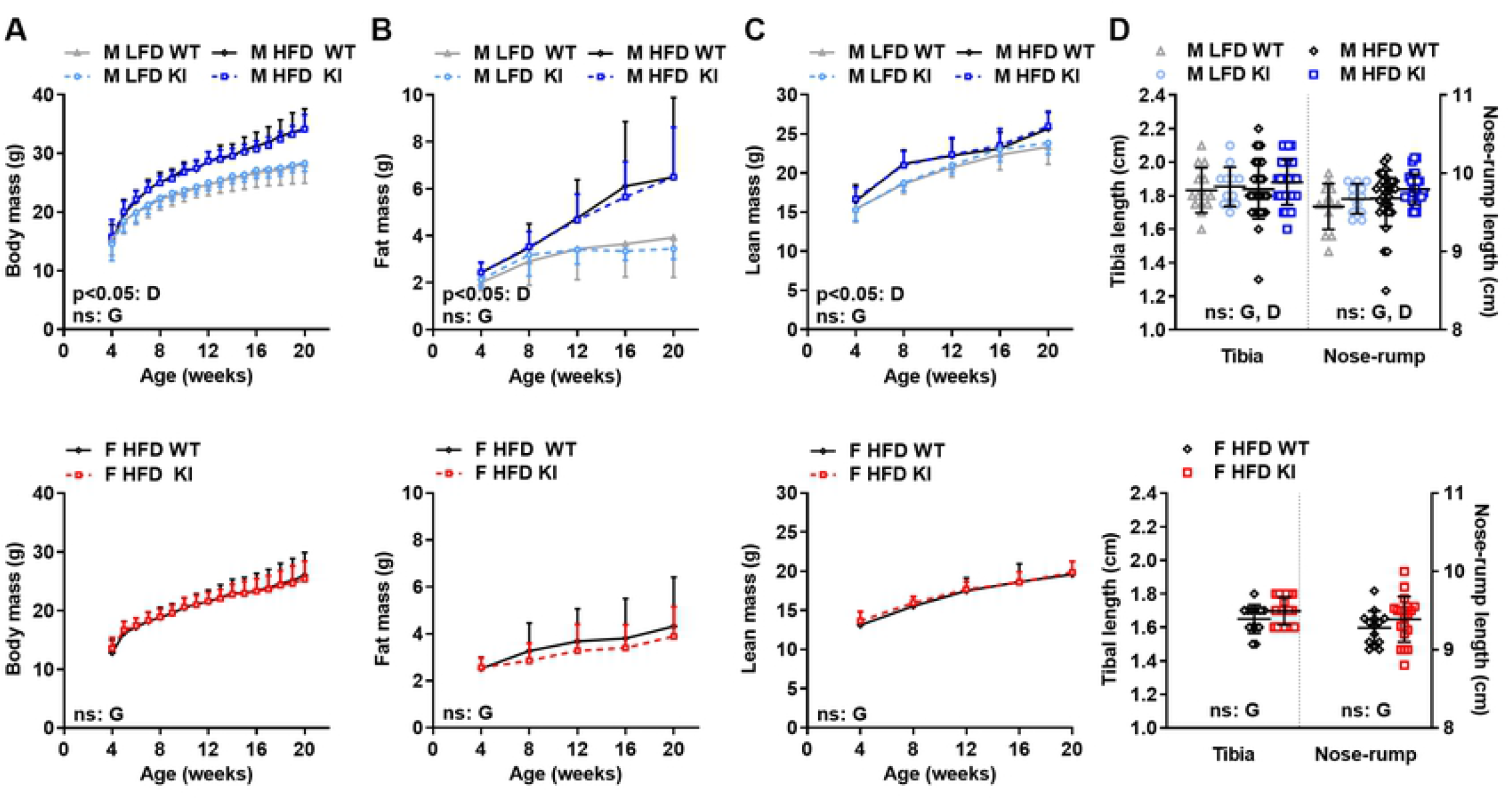
Effect of the CREBRF^R458Q^ variant on body weight, body composition, and linear growth on control diet (low-fat diet) and in response to chronic nutritional overload (high-fat diet) in mice. Male (M, top row, n=12-14/group for LFD and 30-31/group for HFD) and female (F, bottom row, n=27-38/group) wild-type (WT) and knockin (KI) mice were fed low-fat diet (LFD) or high-fat diet (HFD) weaning until sacrifice at 22 weeks of age (or 36 weeks of age in Fig 5). Total body mass (**A**), fat mass (**B**), and lean mass (**C**) were determined at the ages indicated. Tibia and Nose-rump length (**D**) were determined at 22 weeks of age. General linear models with (**A-C**) or without (**D**) time as a within subject variable were used to compare the above dependent variables between D=diet (LFD, HFD) and G=genotype (WT, KI) as fixed factors. Data are mean (SD). Main effects with a significance level of p<0.05 are indicated. Main effects with a significance level of ≥0.05 are labeled as non-significant (ns).

### The CREBRF^R458Q^ variant does not affect energy homeostasis in response to acute (<24h fasting/refeeding) or chronic (high-fat diet) nutritional stress in mice

CREBRF has been shown to be highly regulated by nutritional stress in mammalian cells (6) and Drosophila (18). Since genotype could lead to equal or conditional changes in energy intake and expenditure that do not induce a net change in body mass or composition, we additionally evaluated these parameters in WT and KI mice in response to both acute nutritional stress (<24h fasting and refeeding) and chronic nutritional stress (LFD and HFD feeding). As an initial step, we performed a detailed analysis of phenotypic parameters in association with indirect calorimetry and activity monitoring using a comprehensive laboratory animal metabolic measurement system in 12-week-old male mice fed LFD and HFD. The overall effect for each parameter is expressed as both cumulative values over time (**Fig 3, top row**) and rates per interval (light-dark intervals within fed-fasted-refed intervals) (**Fig 3, bottom row**). No differences were identified between genotypes for total body mass (**Fig 3A**), energy intake per total body mass (**Fig 3B)**, energy expenditure per total body mass (**Fig 3C**), or activity per mouse (**Fig 3A**), regardless of diet, nutritional status, or light cycle. As positive controls, the expected significant effects of diet, nutritional status, and light cycle on these parameters were present, i.e., higher total body mass, energy intake per total body mass, and activity per mouse with HFD compared to LFD, fed/refed compared to fasted states, and dark compared to light cycle; as well as higher energy expenditure per total body mass with fed/refed compared to fasted states, and dark compared to light cycle. Thus, the murine CREBRF^R458Q^ variant does not affect total body mass, energy intake, energy expenditure, or activity in response to either acute nutritional stress (fasting and refeeding) or chronic nutritional stress (LFD and HFD feeding).

**Fig 3.**
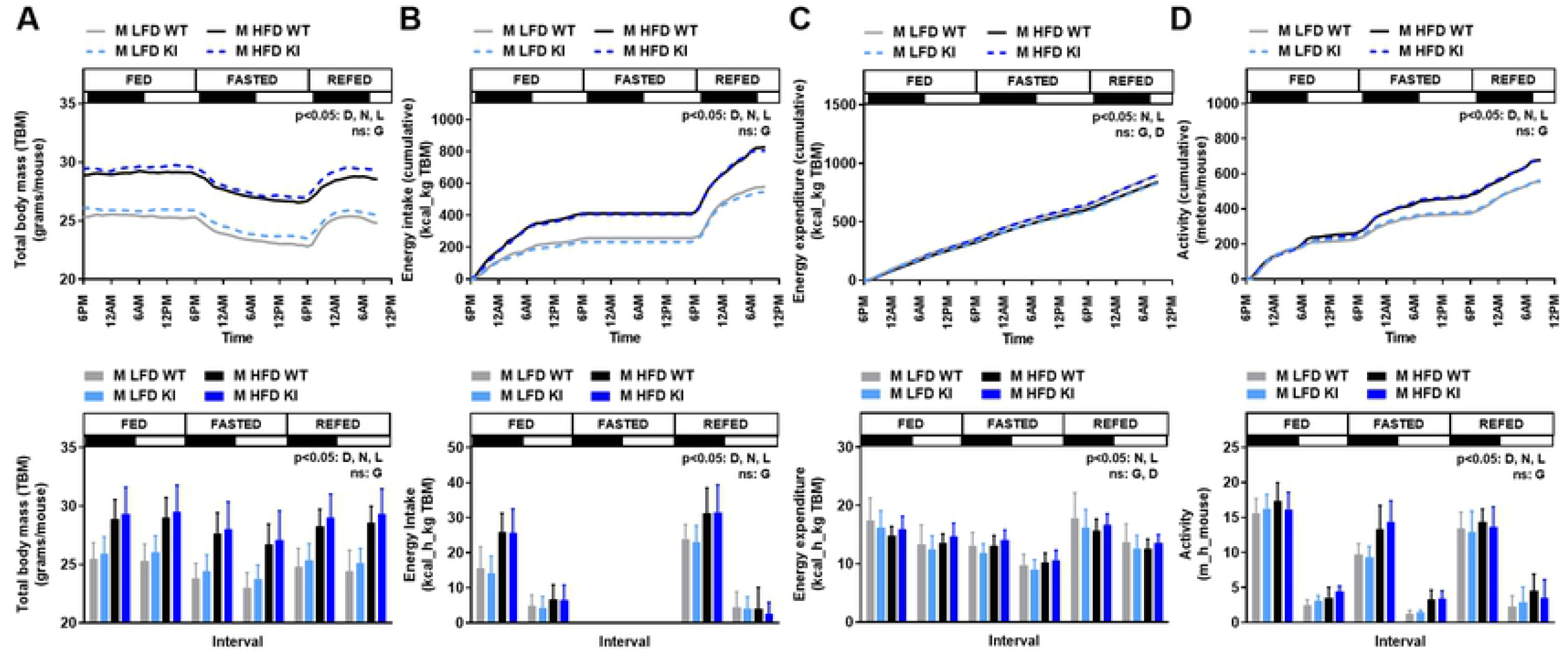
Effect of the CREBRF^R458Q^ variant on energy homeostasis in response to acute (<24h fasting/refeeding) or chronic (high-fat diet) nutritional stress in mice. Male (M, 12-week-old, n=7-14/group) wild-type (WT) and knockin (KI) mice fed control low-fat diet (LFD) or high-fat diet (HFD) were evaluated for changes in total body mass **(A)**, energy intake **(B)**, energy expenditure **(C)**, and activity **(D)** in response to changes in nutritional status (fed, fasted, refed) over time using indirect calorimetry and activity monitors (Sable System). The protocol was as follows: 24-36 hours of acclimation (data not shown), 24 hours of ad lib feeding (6PM-6PM), 24 hours of fasting (6PM-6PM), 15 hours of refeeding (6PM-9AM). Feeding status and lighting status (dark=black bar; light=white bar) are shown on the top of each graph. Data are presented in two ways: Top row = mean (without SD for figure clarity) for each variable over the course of the entire study (**A**=absolute value, **B-D**=cumulative values); or Bottom row = mean (SD) for each variable during each 12-hour light-dark interval (**A**=absolute value; **B-D**=average per hour for that interval). Generalized estimated equations with time as a within subject variable were used to compare the above dependent variables between D=diet (LFD, HFD), N=nutritional status (fed, fasted, refed), L=lighting status (dark, light), and G=genotype (WT, KI) as fixed factors. Main effects with a significance level of p<0.05 are indicated. Main effects with a significance level of ≥ 0.05 are labeled as non-significant (ns).

### The CREBRF^R458Q^ variant does not affect energy substrate utilization, glucose homeostasis, or lipid homeostasis in response to acute (<24h fasting/refeeding) or chronic (high-fat diet) nutritional stress in mice

The human CREBRF^R457Q^ variant is associated with relative protection from metabolic complications of obesity, most notably lower fasting glucose and lower diabetes risk (6, 11-13) as well as absence of expected obesity-associated increase in serum lipids in humans (6). To determine the effect of the murine CREBRF^R458Q^ variant on these parameters, we evaluated energy substrate utilization, glucose homeostasis, and lipid homeostasis in response to acute nutritional stress (<24h fasting and refeeding) and chronic nutritional stress (LFD and HFD feeding) in WT and KI mice. We first determined the respiratory exchange ratio (RER) using the above indirect calorimetry experiment in 12-week-old male mice fed LFD and HFD as above (**Fig 4A**). No differences were identified between genotypes for RER, regardless of diet, nutritional status, or light cycle. As positive controls, the expected significant effects of diet, nutritional status, and light cycle on these parameters were present, i.e., higher RER with LFD-compared to HFD, fed/refed compared to fasted states, and dark compare to light cycle (reflecting greater utilization of carbohydrate over lipid substrates under these conditions). Thus, the murine CREBRF^R458Q^ variant does not affect the ability to switch between glucose and lipid as energy substrates in response to acute or chronic nutritional stress. We next determined blood glucose at 4-week intervals from weaning until sacrifice in HFD-fed male (**Fig 4B-C, blue top**) and female (**Fig 4B-C, red, bottom)** mice in response to fasting (**Fig 4B**) and refeeding (**Fig 4C**). No differences were identified between genotypes for blood glucose at any time point, regardless of nutritional status. As a positive control, the expected significant effects of nutritional status on blood glucose were present, i.e., higher blood glucose in the refed compared to fasted states. We next determined the blood glucose response to an intraperitoneal glucose load using glucose tolerance tests in male (**Fig 4D, blue, top**) and female (**Fig 4D, red, bottom**) mice. No differences were identified between genotypes for glucose tolerance, regardless of diet. As a positive control, the expected significant effects of diet on glucose tolerance were present, i.e., worse glucose tolerance and greater area under the curve for glucose (AUCg) with HFD compared to LFD.

**Fig 4.**
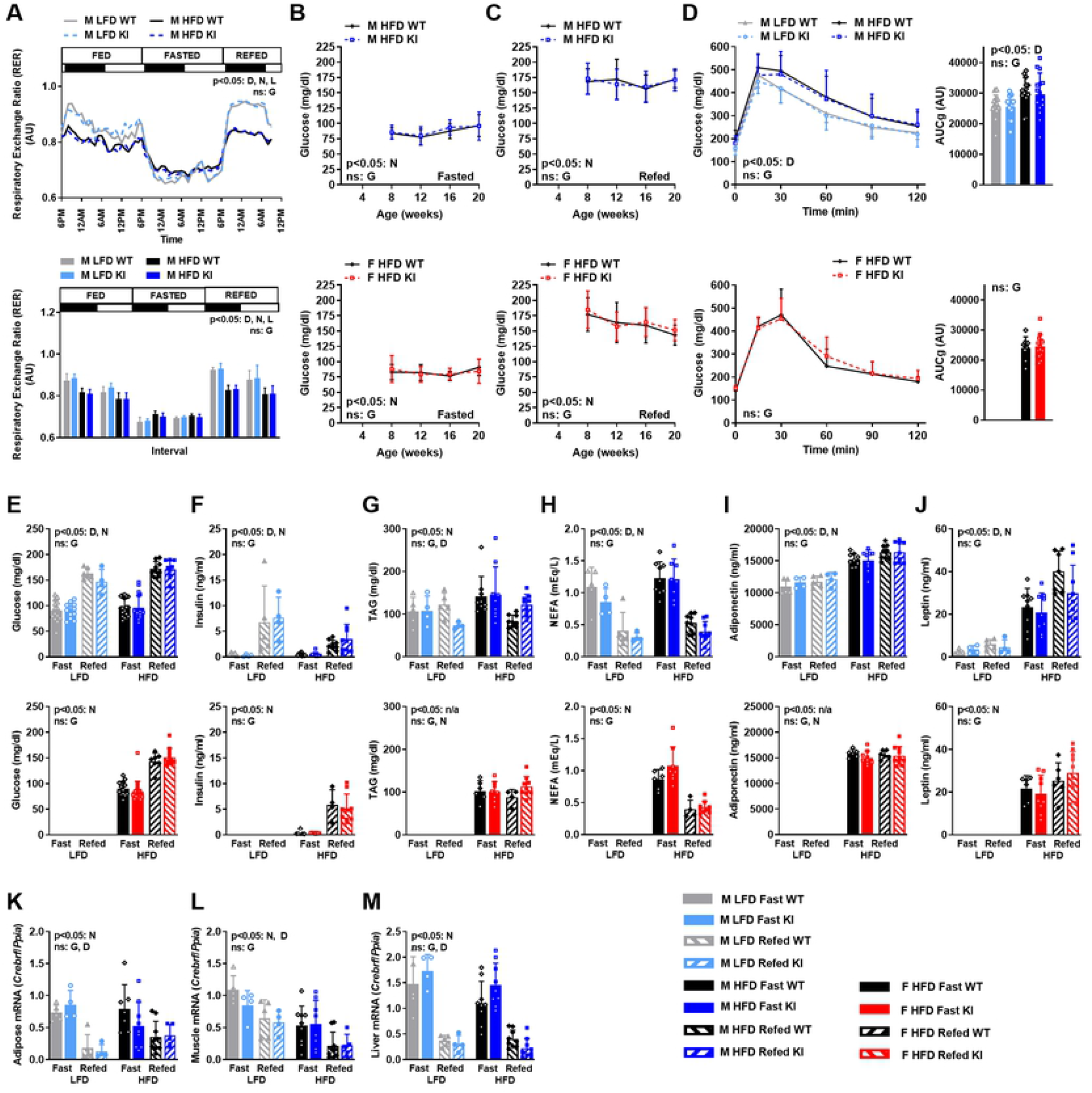
Effect of the CREBRF^R458Q^ variant on energy substrate utilization, glucose homeostasis, and lipid homeostasis in response to acute (<24h fasting/refeeding) or chronic (high-fat diet) nutritional stress in mice. Respiratory exchange ratio (RER) in male (M, 12-week-old, n=7-14/group) wild-type (WT) and knockin (KI) mice fed low-fat diet (LFD) or high-fat diet (HFD) during changes in nutritional status (fed, fasted, refed) over time using indirect calorimetry (Sable System). The protocol, parameters, and analyses are described in the legend for **Fig 3. (B-C)** Blood glucose (dependent variable) at 4-week intervals from weaning until sacrifice in HFD-fed male (M, top, n=18/group) and female (F, bottom, n=14-18/group) wild-type (WT) and knockin (KI) mice in response to fasting (**B**, 16h) and refeeding (**C**, 16h fast followed by 12h refeeding). For clarity, B and C are shown as separate graphs but data are analyzed together. **(D)** Glucose tolerance tests (GTTs, left) and area under the curve for glucose (AUCg, right) in male (M, top, 19-week-old, n=12-18/group) and female (F, bottom, 19 -week-old, n=10-18/group) wild-type (WT) and knockin (KI) mice. **(E-J)** Serum parameters in LFD- and HFD-fed male (E-J, top) and female (E-J, bottom) WT and KI mice sacrificed at 22-weeks-old in the fasted (16h) or refed (24h of fasting followed by 16h of refeeding) states (4-9/group): **(E)** glucose, **(F)** insulin, **(G)** triacylglycerol (TAG), **(H)** non-esterified fatty acids (NEFA), **(I)** adiponectin, and **(J)** leptin. **(K-M)** Expression of *Crebrf* relative to *Ppia* mRNA in adipose tissue **(K)**, gastrocnemius skeletal muscle **(L)**, and liver **(M)** in the same mice as (E-J). General linear models with **(B-D)** or without **(E-M)** time as a within subject variable were used to compare the above dependent variables between D=diet (LFD, HFD), N=nutritional status (fasted, refed), and G=genotype (WT, KI) as fixed factors. Data are mean (SD). Main effects with a significance level of p<0.05 are indicated. Main effects with a significance level of ≥ 0.05 are labeled as non-significant (ns).

We additionally determined serum parameters for glucose and lipid homeostasis in LFD- and HFD-fed male (**Fig 4E-J, blue, top**) and HFD-fed female (**Fig 4E-J, red, bottom**) mice following sacrifice in the fasted and refed states. No differences were identified between genotypes for glucose (**Fig 4E**), insulin (**Fig 4F**), triacylglycerols (**Fig 4G**), non-esterified fatty acids (**Fig 4H**), adiponectin (**Fig 4I**), or leptin (**Fig 4J**), regardless of diet or nutritional status. Likewise, there were no differences between genotypes for hepatic triacylglycerol content under these conditions (data not shown). As a positive control, the expected effects of diet and nutritional status on these parameters were observed, i.e., higher glucose, adiponectin, and leptin with HFD; higher glucose and insulin, lower NEFA and TAG with refeeding. Finally, we measured *Crebrf* mRNA expression and regulation by nutritional status and diet in metabolically-relevant tissues (**Fig 4K-M**), including adipose tissue (**Fig 4K**), skeletal muscle (**Fig 4L**), and liver (**Fig 4M**). *Crebrf* mRNA was regulated by nutritional status (higher with fasting compared to refeeding) in all three tissues and by diet (higher in LFD compared to HFD) in muscle. In contrast, *Crebrf* mRNA expression did not differ by genotype in any of the three tissues. Thus, the CREBRF^R458Q^ variant does not affect energy substrate utilization, glucose homeostasis, or lipid homeostasis in response to acute (<24h fasting/refeeding) or chronic (HFD) nutritional stress in mice, despite regulation of *Crebrf* mRNA expression in response to these interventions.

### The CREBRF^R458Q^ variant does not affect energy or metabolic homeostasis in response to prolonged nutritional stress (>24h fasting) in mice

Evidence from Drosophila (18) and murine 3T3-L1 adipocytes (6) implicates CREBRF in the cellular and physiological adaptation to starvation. We, therefore, hypothesized that physiological manifestations of the murine CREBRF^R458Q^ variant might require the more extreme physiological challenge of prolonged fasting. To test this hypothesis, female WT and KI mice (F, 36-week-old, n=8/group) were fasted until they achieved 25% weight loss, followed by refeeding ad libitum for the same duration, while closely monitoring body mass, fat mass, lean mass, blood glucose, and indirect calorimetry (**Fig 5**) (28). At baseline, littermate weight-matched mice had comparable body mass (**Fig 5A**), lean mass (**Fig 5B**), and fat mass (**Fig 5C**). Mice responded remarkably well to prolonged starvation, exhibiting normal physical and behavioral characteristics. In response to prolonged to fasting, mice lost ∼25% of their initial body mass (**Fig 5A, right**), and the rate of loss with fasting as well as rate of gain with refeeding were similar for percent total body mass (**Fig 5A**), lean mass (**Fig 5B**), and fat mass (**Fig 5C**). Thus, the CREBRF^R458Q^ variant does not differentially affect body mass or composition during prolonged fasting. We next looked at the effect of prolonged fasting on energy substrate homeostasis and utilization. In this smaller sub-cohort, there was a slightly lower baseline blood glucose (p<0.05) and RER (p=0.09) in *Crebrf*KI mice that was inconsistent with result in the larger group (**Fig 4**), likely reflecting a cohort effect in timing since last feeding in the ad lib fed state rather than true difference between groups. Notably, during prolonged fasting, blood glucose (**Fig 5D**) and RER (**Fig 5E**) converged to similar minimums and rebounded identically after refeeding in both genotypes. Thus, the CREBRF^R458Q^ variant does not differentially affect glucose homeostasis or energy substrate utilization during prolonged fasting. Thus, overall, the CREBRF^R458Q^ variant does not affect energy or metabolic homeostasis in response to prolonged nutritional stress (>24h fasting) in mice.

**Fig 5.**
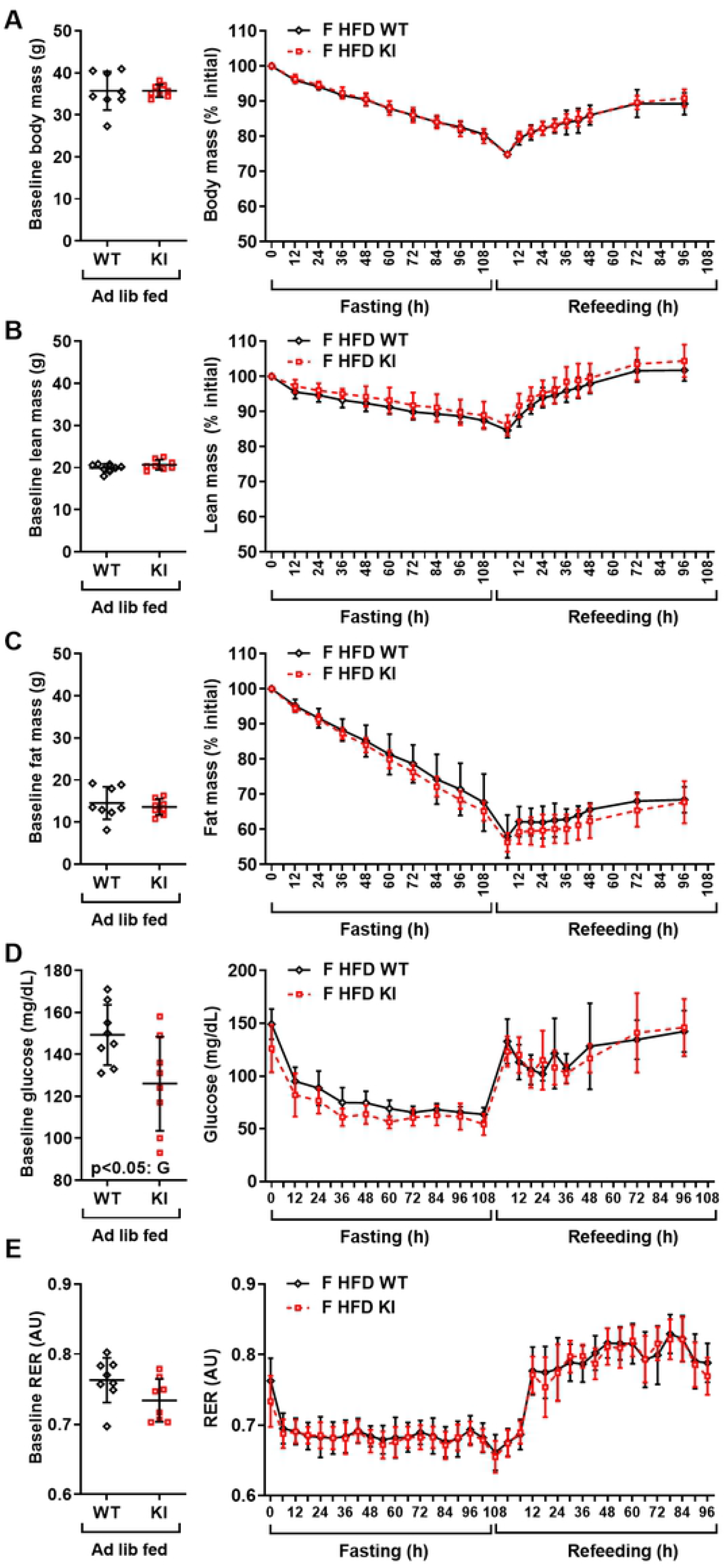
Effect of the CREBRF^R458Q^ variant energy or metabolic homeostasis in response to prolonged nutritional stress (>24h fasting) in mice. Female (F, 36-week-old, HFD-fed, n=8/group) wild-type (WT) and knockin (KI) mice were subjected to a monitored fast until they achieved 25% loss of their initial body mass, followed by refeeding ad libitum. Body mass, lean mass, fat mass, blood glucose, and indirect calorimetry were determined during fasting and refeeding. (**A**) Baseline total body mass after ad lib feeding (**A, left**), and during prolonged fasting and refeeding (**A, right**). (**B**) Baseline lean mass after ad lib feeding (**B, left**), and during prolonged fasting and refeeding (**B, right**). (**C**) Baseline fat mass after ad lib feeding (**C, left**), and during prolonged fasting and refeeding (**C, right**). (**D**) Baseline blood glucose after ad lib feeding (**D, left**), and during prolonged fasting and refeeding (**D, right**). (**E**) Baseline respiratory exchange ratio (RER) after ad lib feeding (E, left), and during prolonged fasting and refeeding (**E, right**). Baseline data (**A-E, left**) were analyzed by unpaired two-tailed t-test. Indirect calorimetry data was analyzed as described in Fig 3. Other data were analyzed using general linear models with time as a within subject variable and G=genotype (WT, KI) as the fixed factor. Data are mean (SD). Main effects with a significance level of p<0.05 are indicated.

### The CREBRF^R458Q^ variant does not affect energy or metabolic homeostasis in response to mTORC1 inhibition in mice

The Drosophila homolog of CREBRF, REPTOR, is induced by the TORC1 inhibitor, rapamycin, and mediates the majority of the downstream transcriptional response to rapamycin-mediated TORC1 inhibition, primarily via its (de)phosphorylation-dependent translocation to the nucleus (18). Although there is no evidence of the latter in mammalian cells, we have confirmed that rapamycin induces CREBRF mRNA expression in murine 3T3-L1 cells (6). We, therefore, hypothesized that the physiological manifestations of the murine CREBRF^R458Q^ variant might be TORC1-specific, and therefore may be unmasked by rapamycin treatment. To test this hypothesis, we treated LFD- and HFD-fed male mice with vehicle or rapamycin 4 mg/kg body weight via intraperitoneal (ip) injection every other day for three weeks prior to sacrifice at ∼22 weeks of age in the ad lib-fed state (**Fig 6**). We first confirmed the effectiveness of our pharmacological intervention. As expected, rapamycin treatment decreased phosphorylated ribosomal S6 (pS6), a known downstream target of TORC1, in all tissues examined (adipose tissue, skeletal muscle, and liver) (**Fig 6A**). We next confirmed the expression and regulation of *Crebrf* mRNA in metabolically-relevant tissues of rapamycin-treated mice (**Fig 6B-D**). Rapamycin increased *Crebrf* mRNA expression in adipose tissue and skeletal muscle but not liver, whereas genotype did not influence *Crebrf* mRNA expression, regardless of treatment. Consistent with previous reports (22), rapamycin treatment resulted in a clear and significant impairment of glucose tolerance and increased area under the curve for glucose (AUCg) in both HFD-fed (**Fig 6E**) and LFD-fed (**Fig 6F**) mice. However, there was no effect of genotype on the glycemic response to rapamycin treatment in mice on either diet. Similar results were obtained for other relevant outcome measures noted in (22) (data not shown). Thus, the CREBRF^R458Q^ variant does not affect energy or metabolic homeostasis in response to mTORC1 inhibition in mice.

**Fig 6.**
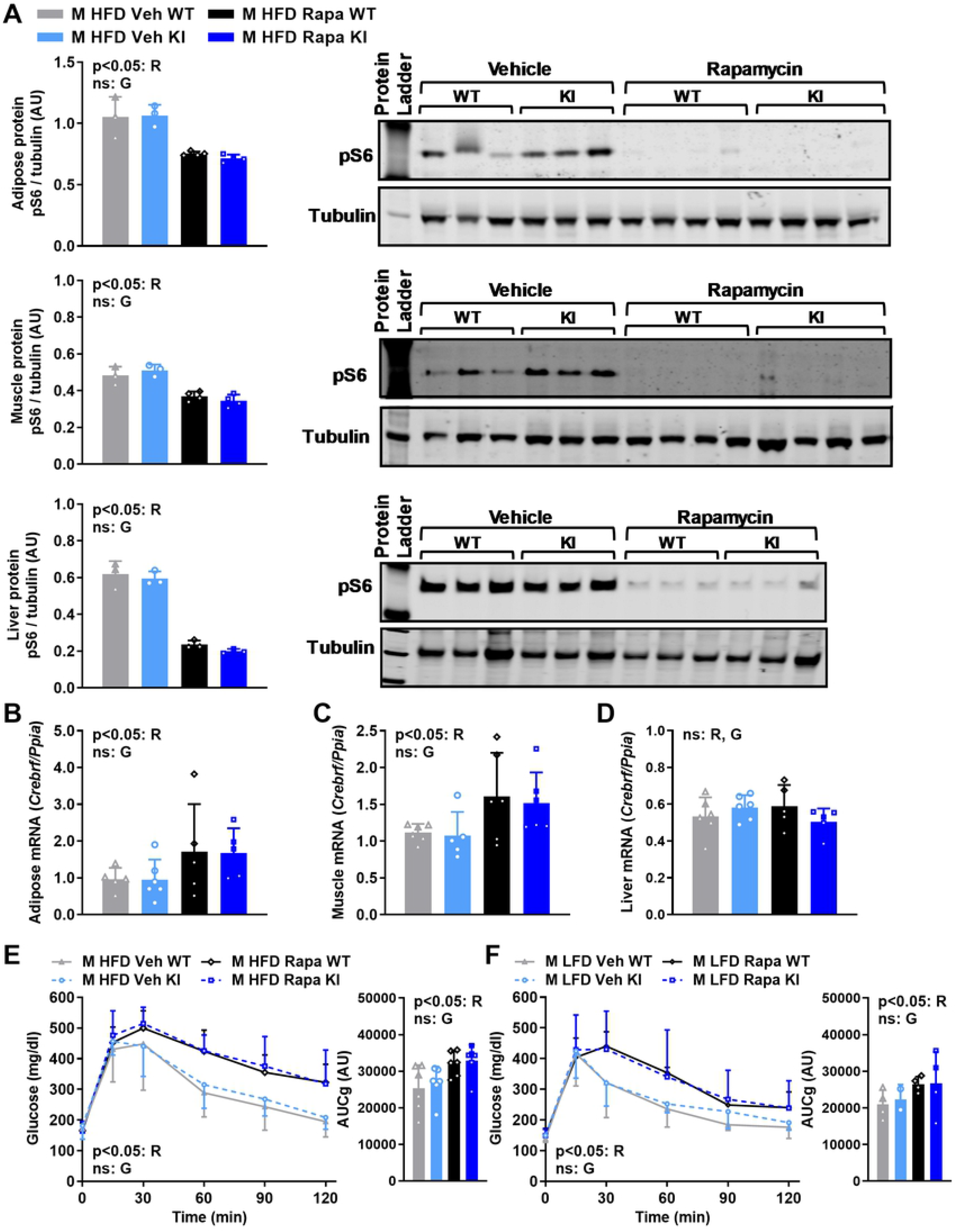
Effect of CREBRF^R458Q^ on energy or metabolic homeostasis in response to mTORC1 inhibition in mice. Male (M, n=6-7/group) wild-type (WT) and knockin (KI) mice were fed low-fat diet (LFD) or high-fat diet (HFD) from weaning until sacrifice at 22 weeks of age were treated with vehicle (Veh) or rapamycin (Rapa) 4 mg/kg body weight via intraperitoneal (ip) injection every other day for three weeks prior to sacrifice at ∼22 weeks of age in the ad lib fed state. **(A)** Protein expression of Ser240/244 phosphorylated ribosomal protein S6 (pS6) relative to tubulin control by Western blotting in adipose tissue (top), gastrocnemius skeletal muscle (middle), and liver (bottom) [left, quantification; right, actual blots). **(B-D)** *Crebrf* mRNA relative to *Ppia* control in adipose tissue **(B)**, gastrocnemius skeletal muscle **(C)**, and liver **(D)** of mice treated with rapamycin. **(E-F)** Glucose tolerance tests (GTTs, left) and area under the curve for glucose (AUCg, right) in rapamycin-treated HFD-fed **(E)** and LFD-fed **(F)** mice. General linear models with **(E-F**, GTTs) or without **(A-D)** time as a within subject variable were used to compare the above dependent variables between R=rapamycin treatment (rapamycin, vehicle) and G=genotype (WT, KI) as fixed factors. Data are mean (SD). Main effects with a significance level of p<0.05 are indicated. Main effects with a significance level of ≥ 0.05 are labeled as non-significant (ns).

## Discussion

The goal of this investigation was to determine the function and physiological relevance of the human CREBRF^R457Q^ obesity/diabetes risk variant by generating and evaluating a murine “knockin” model with the orthologous CREBRF^R458Q^ variant. However, despite strong evidence linking the CREBRF^R457Q^ variant to obesity and diabetes risk in humans, we were not able to detect any genotype effects of the CREBRF^R458Q^ variant on energy, glucose, or metabolic phenotypes in mice. Furthermore, we examined these phenotypes at baseline as well as under conditions of acute (<24h fasting/refeeding), chronic (low-and high-fat diet feeding up to 20w), and extreme (5d starvation) nutritional stress, as well as in response to pharmacological stress (inhibition of TORC1 by rapamycin). Although we did not observe an effect of genotype, we did demonstrate that *Crebrf* mRNA expression is induced by fasting (adipose tissue, skeletal muscle, and liver) and rapamycin-mediated inhibition of TORC1 (adipose tissue, skeletal muscle), which has not been previously reported. The data presented suggest that the CREBRF^R458Q^ knockin mouse model is not likely to be an effective model for understanding the function and physiological relevance of this gene/variant.

There are several possible explanations for why the CREBRF^R458Q^ knockin mouse model did not recapitulate the phenotype of human CREBRF^R457Q^ carriers. Notably, we conducted several quality control experiments to confirm that we indeed successfully engineered the desired orthologous CREBRF^R458Q^ knockin allele in mice. We confirmed this by directly sequencing the allele, as well as by developing three separate genotyping methods and by genotyping mice with at least two methods at weaning and again after sacrifice. Thus, we are confident that we evaluated a valid mouse model. In addition, a separate group independently created a CREBRF^R458Q^ knockin mouse model and likewise found no effect of genotype on body mass, fasting blood glucose, or fasting plasma insulin. Unlike us, they reported an effect on body length in male mice only (10). Importantly, our phenotypic evaluation of the CREBRF^R458Q^ knockin mouse model presented is much more extensive and includes acute, chronic, and extreme nutritional and pharmacological stresses.

It is possible that the CREBRF^R457Q^ variant is causal/relevant in humans, but the orthologous CREBRF^R458Q^ variant in mice is not. A corollary to this possibility is that perhaps the murine CREBRF^R458Q^ variant influences cellular biology in mice that remain undetectable at the physiological level due to compensatory mechanisms. However, we observed no evidence of this. Two other possibilities are that the effect size of CREBRF^R458Q^ variant in mice was too small to detect or was not detectable under the specific conditions tested in mice. To address this issue, we assessed body mass (the primary effect observed in human carriers of this variant) over 22 weeks on all mice generated for this study to maximize power to detect differences. We also focused on the main effects of genotypes for numerous dependent variables at baseline and under nutritional and pharmacological stress. However, we cannot exclude the possibility that an effect of genotype might be detectable under other conditions.

It is also possible that the human CREBRF^R457Q^ variant is not the causative variant contributing to obesity and diabetes risk in humans. However, we do not believe this to be the case for two reasons: 1) the human CREBRF^R457Q^ variant is strongly and repeatedly linked to metabolic traits in humans (especially BMI, glucose, and height), and 2) CREBRF is linked to multiple biological processes known to contribute to these metabolic traits. Regarding the former, in our original report, we confirmed the relationship between the CREBRF^R457Q^ variant and BMI in an initial cohort of 3,072 Samoans and in a separate replication cohort of 2,102 additional Samoans. This relationship between the CREBRF^R457Q^ variant and BMI was highly significant (β 1.356, p 1.12 × 10^−13^), as were the associations with obesity risk (OR 1.305, p 1.12 × 10^−5^) and diabetes risk (OR 0.586, p 6.68 × 10^−9^ after adjusting for BMI) (6). The relationship between CREBRF^R457Q^ and obesity/diabetes risk has now been replicated by numerous groups in other populations across Oceania (9, 11-15). These data provide strong support for the observed associations, but are not sufficient to prove causality.

Regarding the biological evidence linking CREBRF and/or the CREBRF^R457Q^ variant to metabolism, we have previously demonstrated that ectopic overexpression of human CREBRF^R457Q^ in a murine 3T3-L1 adipocyte model promotes adipogenesis, decreases mitochondrial respiration, and increases fat storage (6), consistent with the observed whole-body phenotypes in human carriers of the CREBRF^R457Q^ allele, and providing functional support for a causal role of this variant. Although additional mechanistic studies of CREBRF^R457Q^ variant are still underway, multiple studies have emerged linking (wild-type) CREBRF to multiple proteins/pathways known to be critically involved in energy and metabolic homeostasis including CREB-family proteins (29, 30), TORC1 (18, 31), Akt (32), and glucocorticoid receptor (17, 33) across different species (i.e., human, mouse, goat, Drosophila) and cell types (i.e., adipocyte, neurons, gastric cells, endometrial cells, etc.). In vivo, genetic manipulation of endogenous CREBRF in mice and Drosophila results in dramatic phenotypes related to energy and metabolic homeostasis (17, 18). Specifically, global deletion of CREBRF in mice or Drosophila decreases body mass, whereas the opposite phenotype is observed in human carriers of the CREBRF^R457Q^ allele. We have separately confirmed that global deletion of murine CREBRF in mice decreases body mass and protects against diet-induced obesity, whereas global overexpression of murine CREBRF in mice has the opposite effect (Kershaw, publication in progress). These data suggest that CREBRF does indeed contribute to energy and metabolic homeostasis, and that global and/or tissue-specific targeted deletion and/or overexpression of endogenous murine CREBRF may be more useful mouse models to understand the complexities of this gene. Such studies could lead to more targeted mechanistic hypotheses related to the CREBRF^R457Q^ variant that could be tested using cellular models, especially “knockin” (rather than ectopic expression) models of CREBRF^R457Q^ in human cell lines.

Alternatively, it is possible that one or more variants in linkage disequilibrium with the CREBRF^R457Q^ variant, rather than (or in combination with) the CREBRF^R457Q^ variant itself may be the primary driver(s) of obesity/diabetes risk in humans. In our original report, we actually identified several variants in proximity to CREBRF that were in high linkage disequilibrium (6). We concentrated on three variants: 1) rs12513649 - located 35,727 bp upstream of the CREBRF start site (between the ATP6V0E1 and CREBRF genes), 2) rs150207780 – located in the intronic region between exon1 and exon 2 of CREBRF, 3) rs373863828 (i.e., CREBRF^R457Q^) – located in a highly conserved region (GERP score 5.49) in exon 5 of CREBRF. Since CREBRF^R457Q^ was a missense variant that was predicted to be highly damaging (SIFT, 0.03; PolyPhen-2, 0.996), it was the leading candidate. In addition, a potential causal role is supported by the aforementioned data in murine 3T3-L1 adipocytes. In considering the other variants noted above, both rs12513649 and rs150207780 are both present at minor allele frequencies of 6.8% and 7.7% in East Asian individuals without rs373863828 (5-173045049-C-G, 5-173079668-C-T and 5-173108771-G-A, respectively, in gnomAD 3.1 (34), but no association between them and BMI (35, 36) or T2D (37) has been reported. Though variants far upstream and/or in introns are less likely to contribute to phenotypes, additional studies of these and other variants in linkage disequilibrium with the CREBRF^R457Q^ variant, alone and in combination, may be required to clarify their individual and combined effects in human and mouse cells.

Despite these findings, CREBRF remains a potentially important yet poorly understood gene/protein linked to human health and disease. Thus far, the CREBRF^R457Q^ variant is associated with increased obesity risk and decreased diabetes risk in humans (6, 9, 11-14) as well as height (8, 10) and body composition/lean mass (38). However, the endogenous wild-type (non-variant) CREBRF gene/protein has been linked, either directly or indirectly, to many other clinically-relevant physiological and/or pathological processes in mammals, including susceptibility to viral infection (29), endometrial function during pregnancy (31), angiogenesis (39), neuroendocrine function and behavior (17), Alzheimer’s disease (40), and cancer (32, 41-45). Thus, there continues to be an urgent need to understand the mechanism by which this gene/protein contributes to normal biology and disease, and in particular, the specific effect of the CREBRF^R457Q^ variant, which is highly prevalent in Oceanic populations.

In summary, the CREBRF^R458Q^ knockin mouse model of the human CREBRF^R457Q^ obesity-risk variant does not influence energy or glucose homeostasis in response to nutritional stress. To better understand the physiological relevance of CREBRF, future studies could focus on murine models with global or tissues-specific deletion or overexpression of endogenous CREBRF. Such studies are likely to reveal novel insights into “normal” CREBRF function that would permit more targeted hypothesis testing related to the CREBRF variant in the future. In addition, studies examining the human CREBRF^R457Q^ variant (or variants) in human cells will be instrumental in confirming causality and well as dissecting the underlying mechanisms. Finally, additional studies in human carriers of the CREBRF^R457Q^ variant would enhance the deeper and broader understanding of how this variant might impact human health in the quest to identify tangible strategies for prevention and treatment of obesity and associated metabolic abnormalities in humans.

## Acknowledgements

We acknowledge the University of Pittsburgh Transgenic and Gene Targeting (TGT) Core (Division of Immunology), including Rachael Gordon and Sebastien Gingras for assistance with the design of the targeting and genotyping strategy as well as Chunming Bi for microinjection of zygotes and production of CREBRF^R457Q^ mutant mouse strain. We also acknowledge the University of Pittsburgh Center for Metabolism and Mitochondrial Medicine (C3M) for core researches related to animal phenotyping. Most importantly, we are eternally grateful to our Samoan collaborators at the Samoa Ministry of Health, American Samoa Department of Health and other organizations, as well as the Samoan people for partnering with us to promote knowledge that will ultimately improve the health and well-being of people in Samoa and beyond.

## Author Contributions

**Conceptualization:** Daniel E. Weeks, Stephen T. McGarvey, Ryan L. Minster, Nicola L. Hawley, Erin E. Kershaw

**Data curation:** Jitendra S. Kanshana, Ashlee N. Wood, and Erin E. Kershaw

**Formal analysis:** Jitendra S. Kanshana, Ashlee N. Wood, and Erin E. Kershaw

**Funding acquisition:** Daniel E. Weeks, Stephen T. McGarvey, Ryan L. Minster, Nicola L. Hawley, Erin E. Kershaw

**Investigation:** Jitendra S. Kanshana, Polly E. Mattila, Michael C. Ewing, Ashlee N. Wood, Gabriele Schoiswohl, Anna C. Meyer, Aneta Kowalski, Samantha L. Rosenthal, Erin E. Kershaw

**Methodology:** Sebastien Gingras, Brett A. Kaufman, Ray Lu, Gabriele Schoiswohl, Jitendra S. Kanshana, Erin E. Kershaw

**Project administration:** Jitendra S. Kanshana, Erin E. Kershaw

**Resources:** Sebastien Gingras, Brett A. Kaufman, Ray Lu

**Software:** N/A

**Supervision:** Jitendra S. Kanshana, Erin E. Kershaw

**Validation:** Jitendra S. Kanshana, Erin E. Kershaw

**Visualization:** Jitendra S. Kanshana, Erin E. Kershaw

**Writing – original draft:** Jitendra S. Kanshana, Erin E. Kershaw

**Writing – review & editing:** Jitendra S. Kanshana, Polly E. Mattila, Michael C. Ewing, Ashlee N. Wood, Gabriele Schoiswohl, Anna C. Meyer, Aneta Kowalski, Samantha L. Rosenthal, Sebastien Gingras, Brett A. Kaufman, Ray Lu, Daniel E. Weeks, Stephen T. McGarvey, Ryan L. Minster, Nicola L. Hawley, Erin E. Kershaw

## Supporting information

**S1-S3 Tables. Oligonucleotide (S1 Table), antibody (S2 Table), and key reagent (S3 Table) information**.

**S1 Fig. Comparison of human and mouse CREBRF protein sequence**.

**(A)** Illustration of the amino acid change from an arginine (Arg, R) to a glutamine (Gln, Q) at position 457 in humans (Rs373863828) or 458 in mouse. **(B)** The human and mouse protein sequences are on the top and bottom line, respectively, with the comparison between the two sequences in the middle. Overall, the human and mouse CREBRF protein sequence are highly homologous (∼94%). The human R457 corresponds to mouse R458 (shown in RED).

**S2 Fig. Effect of the CREBRF**^**R458Q**^ **variant on tissue weights at the time of sacrifice in response to nutritional and pharmacological interventions**.

**(A-B)** Tissue weights following acute and chronic nutritional interventions. Male **(A)** and female **(B)** wildtype (WT) and Crebrf^R458Q^ knockin (KI) mice were fed low-fat diet (LFD) or high-fat diet (LFD) from weaning until 22 weeks of age and then sacrificed in the fasted state (16h fast) or refed state (24h fast followed by 16h refeeding) (M, n=7-9 per group; F 7-11 per group). General linear models were used to compare each tissue (dependent variables) between D=diet (LFD, HFD), N=nutritional status (fasted, refed), and G=genotype (WT, KI) as fixed factors. Data are mean (SD). Main effects with a significance level of p<0.05 are indicated.

**S3 Fig. Original Gels**.

